# Seasonal Human Coronaviruses OC43, 229E, and NL63 Induce Cell Surface Modulation of Entry Receptors and Display Host Cell-Specific Viral Replication Kinetics

**DOI:** 10.1101/2023.11.20.567923

**Authors:** Vinayakumar Siragam, Mariam Maltseva, Nicolas Castonguay, Yannick Galipeau, Mrudhula Madapuji Srinivasan, Justino Hernandez Soto, Samar Dankar, Marc-André Langlois

**Author notes:** Authors contributed equally (Vinayakumar Siragam, Mariam Maltseva, Nicolas Castonguay).

## Abstract

The emergence of the COVID-19 pandemic prompted increased interest in seasonal human coronaviruses. 229E, OC43, NL63 and HKU1 are endemic seasonal coronaviruses that cause the common cold and are associated with generally mild respiratory symptoms. In this study, we identified cell lines that exhibited cytopathic effects (CPE) upon infection by three of these coronaviruses and characterized their viral replication kinetics and the effect of infection on host surface receptor expression. We found that NL63 produced CPE in LLC-MK2 cells, while OC43 produced CPE in MRC-5, HCT-8 and WI-38 cell lines, while 229E produced CPE in MRC-5 and WI-38 by day 3 post-infection. We observed a sharp increase in nucleocapsid and spike viral RNA (vRNA) from day 3 to day 5 post-infection for all viruses, however the abundance and the proportion of vRNAs copies measured in the supernatants and cell lysates of infected cells varied considerably depending on the virus-host cell pair. Importantly, we observed modulation of coronavirus entry and attachment receptors upon infection. Infection with 229E and OC43 led to a downregulation of CD13 and GD3, respectively. In contrast, infection with NL63, and also with OC43, lead to an increase in ACE2 expression. Attempts to block entry of NL63 using either soluble ACE2 or anti-ACE2 monoclonal antibodies demonstrated the potential of these strategies to greatly reduce infection. Overall, our results enable a better understanding of seasonal coronaviruses infection kinetics in permissive cell lines, and reveal entry receptor modulation that may have implications in facilitating co-infections with multiple coronaviruses in humans.

**IMPORTANCE:** Seasonal human coronavirus are an important cause of the common cold associated with generally mild upper respiratory tract infections that can result in respiratory complications for some individuals. There are no vaccines available for these viruses, with only limited antiviral therapeutic options to treat the most severe cases. A better understanding of how these viruses interact with host cells is essential to identify new strategies to prevent infection-related complications. By analyzing viral replication kinetics in different permissive cell lines, we find that cell-dependent host factors influence how viral genes are expressed and virus particles released. We also analyzed entry receptor expression on infected cells and found that these can be up or down modulated depending on the infecting coronavirus. Our findings raise concerns over the possibility of infection enhancement upon co-infection by some coronaviruses, which may facilitate genetic recombination and the emergence of new variants and strains.

## INTRODUCTION

Human coronaviruses (HCoVs) OC43, 229E, NL63 and HKU1 are seasonal endemic viruses circulating worldwide that cause mostly non-severe upper tract respiratory infections (1–5). It is estimated that they account for 15% to 30% of all the common cold case (6). Epidemiological studies on respiratory infections have reported that OC43 is the most prevalent strain in humans followed by NL63, 229E and HKU1 (7–9). While HCoVs generally cause mild symptoms in healthy individuals, some reports have shown that NL63 and 229E can cause life-threating respiratory problems such as pneumonia leading to respiratory distress syndrome in immunocompromised individuals (10, 11).

Phylogenetically, OC43 and HKU1 belongs to the betacoronavirus genus that includes highly pathogenic coronaviruses such as MERS-CoV, SARS-CoV, and SARS-CoV-2. Coronaviruses 229E and NL63, both belong to the alphacoronavirus genus (12). OC43, 229E and NL63 use different viral attachment and entry receptors including peptidases, cell adhesion molecules and sugars. In the case of 229E, the cellular receptor is CD13 (aminopeptidase N), while OC43 and HKU1 both bind sialic acid (9-O-acetylated ganglioside; GD3) for cell attachment, and HKU1 uses the protease TMPRSS2 as its protein receptor (12–15). NL63 makes use of heparin sulfate for cell attachment and ACE2 for cell entry, while the protein receptor for OC43 has still not been identified (16, 17). Of interest, previous studies have reported the inhibitory role of soluble ACE2 (sACE2) to prevent SARS-CoV-2 infection, which also shares ACE2 as an entry receptor with NL63 (18). However, the blocking effect of sACE2 or monoclonal ACE2 antibody therapy for NL63 infections have not been reported. Alternatively, interferons as potential antiviral regulators against OC43 and 229E infections have been recently studied (19). Although some antiviral drugs like the RNA-dependent RNA polymerase inhibitor remdesivir and the protease inhibitor nirmatrelvir can inhibit OC43, 229E and NL63 in laboratory assays (20–22), these are not commonly prescribed to infected patients. In fact, there are very few antiviral drugs and no vaccines for common cold coronaviruses (23).

Although seasonal coronaviruses have been studied using animal models (24), most of our knowledge of these viruses has been acquired using cell line models (4, 25–27). Cell lines of various types and animal origins have been used to understand the mechanism of virus entry into cells (15, 17, 28), viral replication (4, 29), cytopathic effects (26, 30) and to test the neutralizing ability of antibodies (31). While viral replication and pathogenesis have been extensively characterized for highly pathogenic coronaviruses like SARS-CoV-2, there is a gap in knowledge of HCoV viral replication kinetics in permissive cell lines and their effects on the host cells. In this study, we identified permissive cell lines susceptible to cytopathic effects (CPE) upon OC43, 229E, and NL63 infection and then characterized replication kinetics in these cells by tracking vRNA transcripts coding for the nucleocapsid (N) and spike (S) proteins. We also investigated the modulation of key cell receptors and attachment markers following infection. Finally, we tested the effects of soluble ACE2 and anti-ACE2 monoclonal antibodies at blocking NL63 infection. This detailed study of HCoV infection provides a better understanding of the replication and infection of these viruses in cell culture models which can help prompt the development of treatments, antivirals and cures for the common cold.

## MATERIALS AND METHODS

### Cell culture

HCT-8 cells (human ileocecal adenocarcinoma; ATCC/CCL-244), HEK 293T cells (human embryonic kidney; CRL-3216/ATCC); HEK293/ACE2-stable cell line (HEK293 cells stably expressing angiotensin-converting enzyme 2 (ACE2); BEI Resources/NR-52511) were grown in RPMI-1640 medium supplemented with 100 units/ml penicillin (P), 100 μg/ml streptomycin, and 10% FBS. MRC-5 cells (human lung fibroblasts; CCL-17/ATCC), WI-38 cells (human fetal lung; CCL-75/ATCC), Vero E6 cells (African green monkey kidney cells; CRL-1586/ATCC) and Calu-3 cells (human non-small-cell lung adenocarcinoma; HTB-55/ATCC) were grown in DMEM (MUTICELL#319-005-CL) medium supplemented with 100 units/ml penicillin, 100 μg/ml streptomycin, and 10% FBS. BEAS-2B cells (human bronchial epithelial; CRL-9609ATCC) cells were grown in bronchial epithelial cell growth basal media (BEBM; Lonza/CC-3170). LLC-MK-2 cells (rhesus monkey kidney; CCL-7/ATCC) were cultured in MEM (GIBCO #11095-080) supplemented with 100 units/ml penicillin and 100 μg/ml streptomycin (P/S), 2 mM L-glutamine, 1x non-essential amino acids (NEAA), and 3% FBS. All cell lines were cultured in a ventilate T-75 flask at 37°C in a 5% CO2 humidified incubator. All cell lines were tested for mycoplasma contamination and remained negative.

### Seasonal virus strains and stock propagation

Virus strains were obtained from ATCC and were used to infect their recommended permissive cell line. 229E (stock, VR-740/ATCC) was propagated on MRC-5 cells, OC43(CR-1558/ATCC) on HCT-8 cells and NL63 (BEI Resources/NR-470) on LLC-MK2 cells. The permissive host cells (MRC-5, or HCT and LLC MK2) were maintained at a confluency of 80-90% 24 to 48 hours prior to infection. The cells were infected with a MOI of 0.1, or 0.01 at 35°C for 229E, 33°C for OC43 and 34°C for NL63 for one hour for virus adsorption with rocking every 10 to 20 minutes. The virus culture media for 229E (DMEM +3% FBS+ 1% P/S), OC43 (RPMI +3% FBS +1% P/S) and NL63 (MEM + 3% FBS +1% P/S) was added to the dish (e.g. 1 ml per 25 cm2) to stop the virus adsorption. Viruses were harvested by scrapping the infected cells with a cell scrapper followed by centrifugation at 500 xg for 10 minutes. The supernatant was collected in aliquots and quickly frozen at -80 °C.

### TCID50 assay and CPE

Virus titers of 229E, or OC43 and NL63 were determined by tissue culture infection dose 50 (TCID_50_) assay on their ATCC-recommended cell lines. MRC-5 (20,000 cells), HCT-8 cells (100, 000 cells) and LLC-MK2 cells (300, 000 cells) were seeded at their respective densities in 48 well plate **(Fig. 1).** The cells were checked for confluency (70% to 80%) and the virus dilutions (10-fold serial dilution) were performed. Cell culture media (360 µl) was added into each well and 40 µl of the virus dilution (10-fold) was added to each well. Mock represents the negative control having media only. The plates were incubated at their respective temperatures (229E at 35°C; OC43 at 33°C; NL63 at 34 °C/5% CO2). The plates were shaken every 20 minutes. After 1 to 2 hours of incubation, media was removed, and 400 µl of the culture media supplemented with 1x DMEM-high glucose + 10% BSA +100x NEAA + 100x (100mM) Sodium Pyruvate + 100x (200mM) L-glutamine, 100x Penicillin (10,000 Units/ml)-Streptomycin (10,000 µg/ml) and TPCK-treated trypsin (0.5 µg/ml) was added. TCID_50_ of the viruses were calculated according to Reed–Muench method (32). Based on virus titers (TCID_50_) and cell number, MOI (multiplicities of infection) were determined for the infection experiments

**Figure 1.**
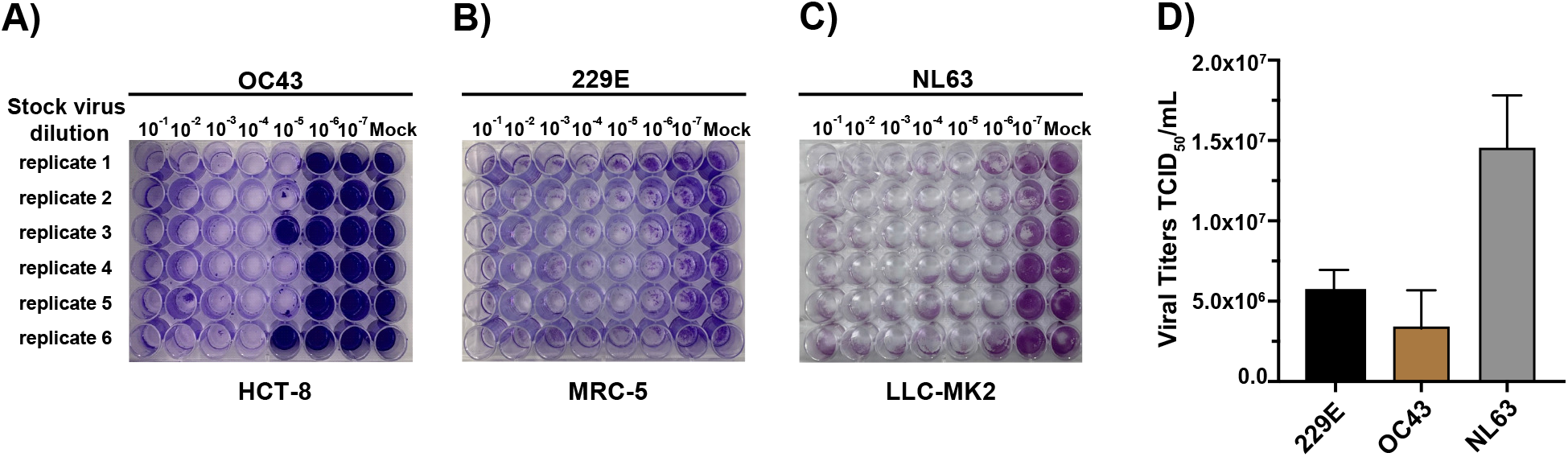
OC43, 229E and NL63 titration by TCID_50_. (A) HCT-8 cells, (B) MRC-5 cells and (C) LLC-MK2 were grown in 48-well tissue culture plates under standard conditions. When the cells (A-C) were confluent (75%), the media was removed, and the cells were infected with seven different 10-fold dilutions (10^-1^ to 10^-7^) of OC43 virus (HCT-8), 229E virus (MRC-5) or NL63 virus (LLC-MK2). The cytopathic effect (CPE) in the wells was monitored for 7 days under a ZOE Cell Imager. A significant CPE is noted as cell death and disturbed cell morphology. The number of wells with CPE (empty wells) and without CPE (filled with crystal violet stain) were counted according to Reed and Muench method (31). (D) Shows the calculation of viral TCID_50_ titers.

All cell lines detailed in **Table 1** were tested for cytopathic effects (CPE) following infection at an MOI of 0.1 by all three HCoVs studied here. The cells were observed daily for CPE following infection under a ZOE Cell Imager (BIO-RAD/1450031) using its phase-contrast setting to monitor cell rounding, clumping and detachment 3 days post infection (dpi). CPE was rated empirically by operator perceived intensity of cellular deformation and detachment: +: less than 20% cell detachment and mild morphological changes start to appear; ++: 20-60% cell detachment and clear morphological changes in most cells; +++: 60-100% cell detachment and severe morphological changes.

**Table 1.** Susceptibility of various cell lines to CPE following infection by HCoVs.

| Cell line tested | Cell type | Human coronavirus (HCoV) tested | Intensity of CPE at day 3 pi* |
| --- | --- | --- | --- |
| BEAS-2B | Human bronchial | OC43 | - |
|  |  | 229E | - |
|  |  | NL63 | - |
| Calu-3 | Human lung cancer | OC43 | - |
|  |  | 229E | - |
|  |  | NL63 | - |
| HCT-8 | Human colon cancer | OC43 | + |
|  |  | 229E | - |
|  |  | NL63 | - |
| LLC-MK2 | Rhesus monkey kidney cells | OC43 | - |
|  |  | 229E | - |
|  |  | NL63 | ++ |
| MRC-5 | Human fetal lung | OC43 | +++ |
|  |  | 229E | + |
|  |  | NL63 | - |
| Vero E6 | African green monkey kidney | OC43 | -/+** |
|  |  | 229E | - |
|  |  | NL63 | - |
| WI-38 | Human fetal lung | OC43 | +++ |
|  |  | 229E | ++ |
|  |  | NL63 | - |
| 293T and 293T ACE2 | Human embryonic kidney | OC43 | - |
|  |  | 229E | - |
|  |  | NL63 | - |

\* Infection was confirmed by detection of both S and N by ddPCR. Infection was performed at a MOI of 0.01. CPE was assessed by phase-contrast light microscopy. +: less than 20% cell detachment and mild morphological changes start to appear; ++: 20-60% cell detachment and clear morphological changes in most cells; +++: 60-100% cell detachment and severe morphological changes. \*\* Very mild CPE at 3dpi was inconsistent across biological replicates (n =3).

### HCoV infection and RNA extraction

MRC-5, HCT-8 and LLC-MK2 cell lines were cultured to reach 60 to 80% confluence in 6-well plates. The following day cell lines were inoculated with their respective virus strains with at MOI 0.01 and 0.1. Virus adsorption was done for 1 hour at 35°C (229E virus), 33°C (OC43 virus) or 34 °C (NL63) followed by the addition of the respective virus culture media. Cells were infected with their respective virus strains and further incubated for the following days to monitor the infection and CPE **(Table 1**): 7 days for OC43, 5 to 7 days for 229E and 7 to 10 days for NL63. HCoV-infected cells were examined under ZOE Cell Imager for the CPE formation followed by total vRNA extraction (QIAamp vRNA Mini kit (QIAGEN # 52904) from culture supernatants and cell lysates along with uninfected cells (control mock) on different dpi (1 to 5). Absolute quantification of vRNA copies of N and S for OC43, 229E or NL63 viruses were determined by Bio-Rad One-Step Reverse Transcriptional Droplet digital PCR (RT-ddPCR).

### RT-ddPCR on vRNA

Duplex, probe-based, one-step RT-ddPCR (1-Step RT-ddPCR Advanced Kit for Probes, Bio-Rad, Hercules, CA) was used to determine absolute quantification of N and S vRNA copy numbers in culture supernatants and cell lysates using the OC43, 229E or NL63 primer-probe sets and 18S was used as reference gene control to normalize the levels of intracellular vRNA. Primers and probes used in this study were synthesized by Integrated DNA Technologies, Inc (IDT, Kanata, Canada) **(Table 2)**. For all the experimental samples, a master mix for each virus (e.g., OC43, or 229E, or NL63) was prepared having 5.5 µl One-Step RT-ddPCR advanced supermix, 2.2 µl RT, 1.1 µl DTT, 1.1 µl (250 nM) 20x primer/probe, 5.5 µl RNA template and 6.6 µl RNase-free water to make a total reaction volume of 22 µl. Blank is considered as nuclease free water as the RNA extraction control. Mock uninfected and infected samples were included with three biological replicates. The reactions were then emulsified by using a QX200 droplet generator (Bio-Rad, Hercules, CA). Droplets that are generated were transferred to a new microplate, and PCR was performed in a thermocycler C1000 (Bio-Rad, Hercules, CA). The reaction conditions for target gene N and S were as follows: reverse transcription (RT) step was performed at 50°C, for 60 min, followed by polymerase activation at 95°C/10 min, and 39 cycles of denaturation at 95°C/30 s, annealing/extension at 55°C/60 s (target gene N), or 60°C/60 s (target gene S), respectively. Deactivation of polymerase was done at 98°C for 10 minutes and droplets were stabilized at 4°C for 30 minutes. Droplets were then read by an Bio-Rad two channel automated QX200 droplet reader. The results were then analyzed by Quanta soft Analysis Pro v.1.0 (Bio-Rad, Hercules, CA). VRNA levels of target genes N or S were normalized to 18S. In this article, the data from the ddPCR experiments are presented as vRNA genome copies per µl (VRG) of reaction mix calculated from the number of positive and negative droplets in the samples based on the following equation (33). VRG = (C.V1)/(5.V2) x D; C = number of copies per 20 µl of reaction mix; V1 = elution volume of extracted RNA in µl (e.g., 60 µl); V2 = sample volume used for RNA extraction in µl (e.g., 140 µl) and D = dilution factor (df) for vRNA. The following dilutions were applied int the above equation for RNA viruses-NL63 (df: 10,000), OC43 (df: 5000) and 229E (df: 50,000).

**Table 2.**
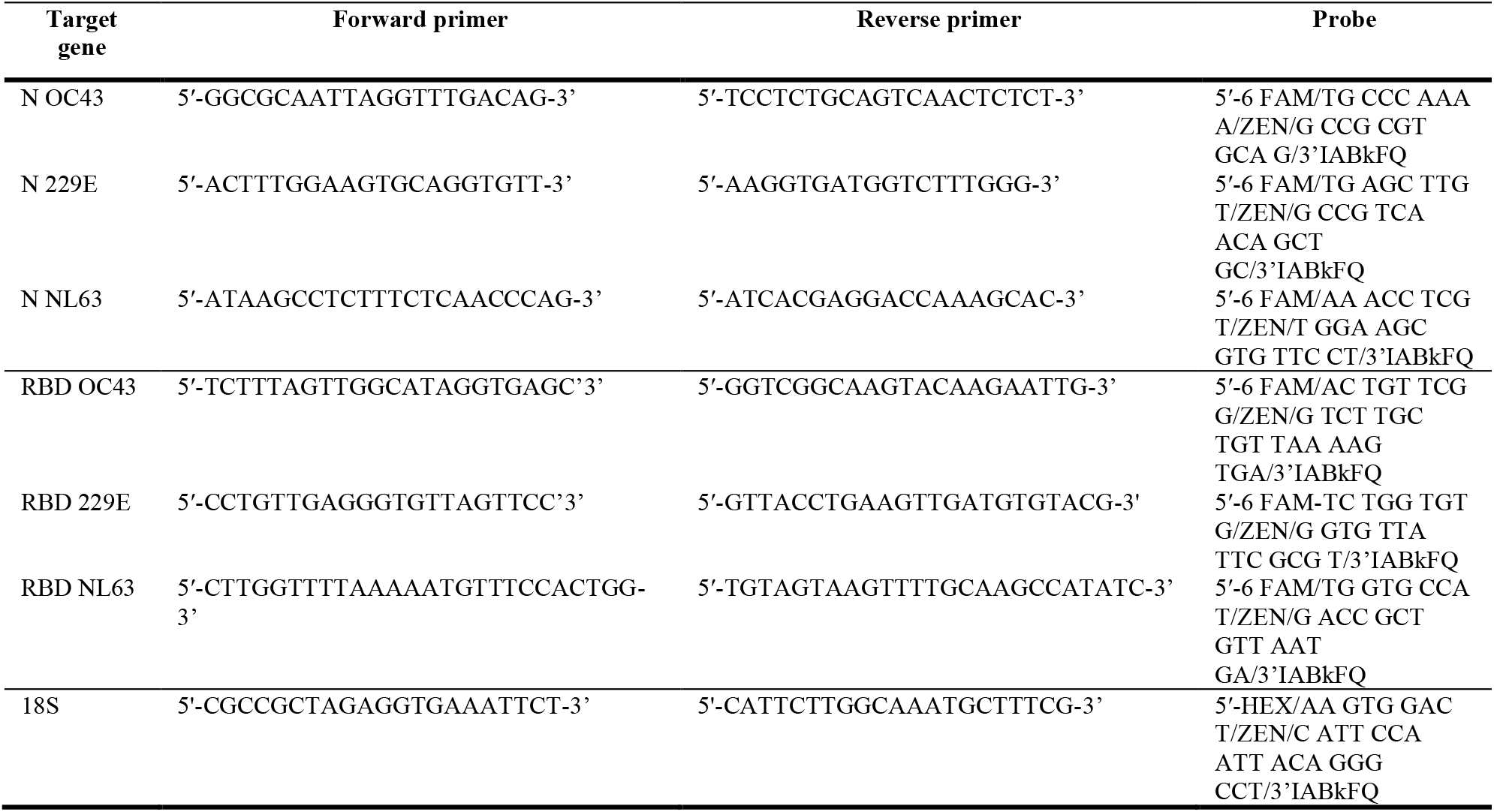
Primers and probes used for ddPCR.

### Detection of surface markers and receptors by flow cytometry following coronavirus infection

Permissive cells (MRC-5, HCT-8 and NL63) were cultured in 6-well plate to attain 75% confluency. Cells were infected at a MOI of 0.1. Five days post-infection, cells were harvested to prepare single cell suspension by lifting the cells with 5 mM EDTA followed by centrifugation at 500 x *g* for 5 minutes and 0.5 ×10^6^ cells were aliquoted per FACS tube. The following antibodies and isotypes were used to quantify the surface marker expression and receptor modulation in the uninfected (control) and infected cells. PE anti-human CD13, anti-human CD147, (isotype-control antibody mouse IgG1 κ, BioLegend); and anti-human CD326 (isotype-control antibody mouse IgG2a κ; BioLegend); Human anti-ACE2 antibody (isotype-control antibody Goat IgG; R&D Systems) and anti-GD3, 9-O-acetyl (clone UM4D4; isotype-control antibody mouse IgM κ; Millipore Sigma). Cells were stained with CD13, CD147, CD326, ACE2 and GD3 antibodies and their corresponding isotypes were incubated for 30 min on ice followed by washing with 0.2% BSA-PBS two times. Cells stained with anti-ACE2 and anti-GD3 were further incubated with secondary antibodies Alexa Fluor 488-labeled sheep anti-goat and goat anti-mouse (Invitrogen) for 30 minutes. After incubation cells were washed twice with 0.2% BSA-PBS, resuspended in 2% BSB-PFA and analysed by Flow Cytometry on BD Celesta instrument (Beckman Coulter) using FlowJo^TM^ software. Flow cytometry data was further analyzed using FlowJo Software v.10.7.1 (FlowJo, Ashland, OR). For cell analysis, SSC-area vs. FSC-area followed by FSC-height vs. FSC-area and SSC-area vs. FSC-area were used to exclude cell aggregates and debris. Live/dead stain BV421^TM^dye was used to excluded dead cells from analysis.

### Blocking of NL63 infection by soluble ACE2 and monoclonal anti-ACE2 antibody

LLC MK2 cells were seeded in 48 wells plates (5×10^5^ cells per well) in MEM medium containing 3% FBS. Prior to incubation of soluble ACE2 (sACE2, National Research Council Canada) with NL63 virus on LLC MK2 cells, the following dilution of sACE2 and NL63 virus was prepared. sACE2 (5 mg or 15 mg), or control protein OVA albumin (Sigma/A5503) were mixed with NL63 virus at MOI 0.1 (1:1) in a final volume of 100 µl in a separate 48 well plate in MEM medium (3% FBS) at 37°C for 30 min. After incubation, the sACE2 + NL63 virus mixture were added to the host cells LLC MK2 (24 hrs post-seeding) and incubated at 34°C for a second incubation for 24 hrs. The following day, the treated LLC MK2 cells were washed with PBS and replaced with infection media (MEM 3% FBS) and cells were monitored to check the blocking effect on NL63 infection until 7 dpi. At the end point day 7, the culture supernatants and cell lysates are harvested to measure the vRNA copies of N and S by ddPCR following vRNA extraction. In a similar approach to monitor the blocking effect on NL63 infection using anti-ACE2 (human), monoclonal blocking antibody that recognizes ACE2 (AdipoGen Life Sciences#AC384), host LLC MK2 cells (48 well plate, 5×10^5^ cells/well) were incubated with human ACE2 monoclonal blocking antibody (10, 40 and 120 µg/ml), or unrelated control antibody (40 µg/ml; anti-DC-SIGN) for 1 to 2 hrs with intermittent shaking followed by the addition of NL63 virus (MOI # 0.1) in a final volume of 100 µl per well at 34°C. After 24 hrs post infection, cells were washed with PBS and replaced with infection media (3%).

### Statistical analysis

Statistical analyses were performed with multiple unpaired t-test with Welch’s correction and unpaired Mann-Whitney test was performed using GraphPad Prism (GraphPad Software, San Diego, CA version 9.5.1).

## RESULTS

### Characterization of cell lines and evaluation of cytopathic effects

We selected a number of cell lines that were previously reported in the literature to be permissive to infection and viral replication of HCoVs (**Table 1**) (4, 25, 27, 29, 34). Our first objective was to identify the cell lines most susceptible to infection by HCoV and that also display cytopathic effects (CPE) within a short 3-day time frame. We infected these cells with the three most prevalent HCoVs, OC43, 229E and NL63, and assessed which ones displayed CPE that resulted in morphological changes and dysmorphic appearances in the cell monolayer with disturbed cell-to-cell aggregates or clumps after 3 days. The intensity of CPE, ranging from - to +++, was determined empirically by phase-contract microscopy. Cells exhibiting CPE were tested by ddPCR to confirm the expression of S and N vRNAs. We used a TCID50 assay (**Fig. 1)** to determine the virus titres for infection by taking into consideration the number of wells with and without CPE based on the methodology of Reed and Muench (32). Permissive cell lines identified in this study are HCT-8 (human colon cancer), MRC-5 (human fetal lung), LLC-MK2 (Rhesus monkey kidney) and WI-38 (human fetal lung). Infection of HCT-8, WI-38 and MRC-5 with OC43 virus showed apparent CPE at day 3 pi (post-infection) with complete disturbance (+++) of the cell monolayer at day 5 pi **(Fig. 2A, 2D and 2G**). MRC-5 and WI-38 cells infected with 229E showed increasing CPE from day 3 pi (**Fig. 3A and 3D)**. Similarly, LLC-MK2 cells also started to display CPE upon NL63 infection from day 3 pi **(Fig. 4A)**. At day 5 pi, the infected LLC-MK2 cells showed aggregation. We also assessed infection of MRC-5, WI-38 and HCT-8 cells infected with NL63, however no CPE or viral replication was measured in these cells, in agreement with the previous reports (25).

**Figure 2.**
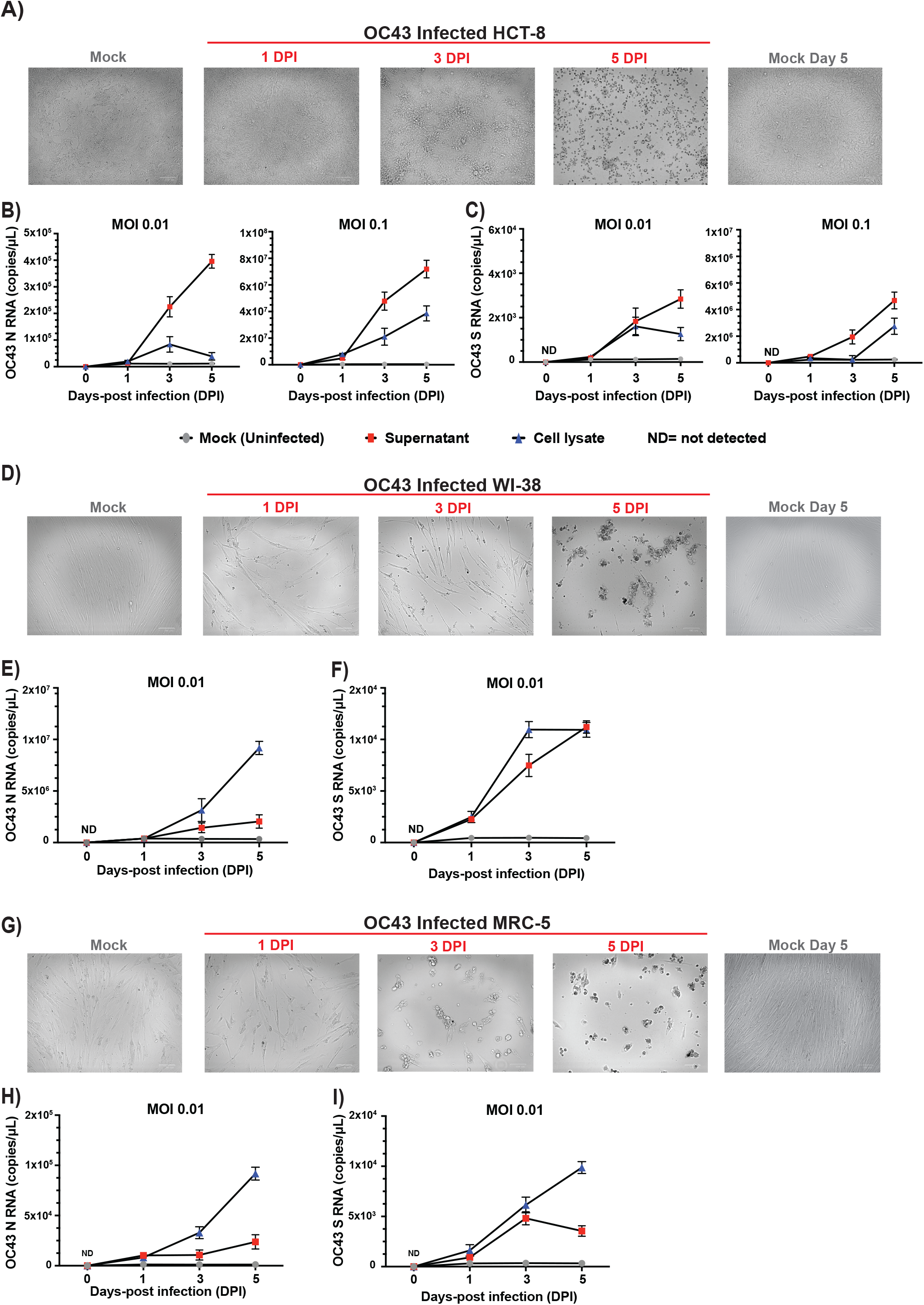
CPE evaluation and replication kinetics of OC43 infection. (A) HCT-8, (D) WI-38 or (G) MRC-5 cells were infected with OC43 (virus titer: 8.89 ×10^5^ TCID50/ml) at a MOI of 0.01. CPE of the infected cells along with mock (uninfected cells) were monitored for morphological changes in the cell monolayer. Images of the cells were captured at different days post infection (day 0 to day 5 dpi) along with uninfected mock by ZOE Cell Imager. The white reference bar is 100 microns. HCT-8 cells (B and C), WI-38 cells (E and F) or MRC-5 cells (H and I) infected with OC43 virus (MOI= 0.01 and/or 0.1) and cell lysates and supernatants were harvested at 0, 1, 3 and 5 days post-infection (dpi). Viral RNA copies of nucleocapsid (N), panels B, E and H, and spike (S), panels C, F and I, were then quantified by ddPCR. ND; not detected. Data represents 3 biological replicates with 3 technical replicates per time point (n = 9).

**Figure 3.**
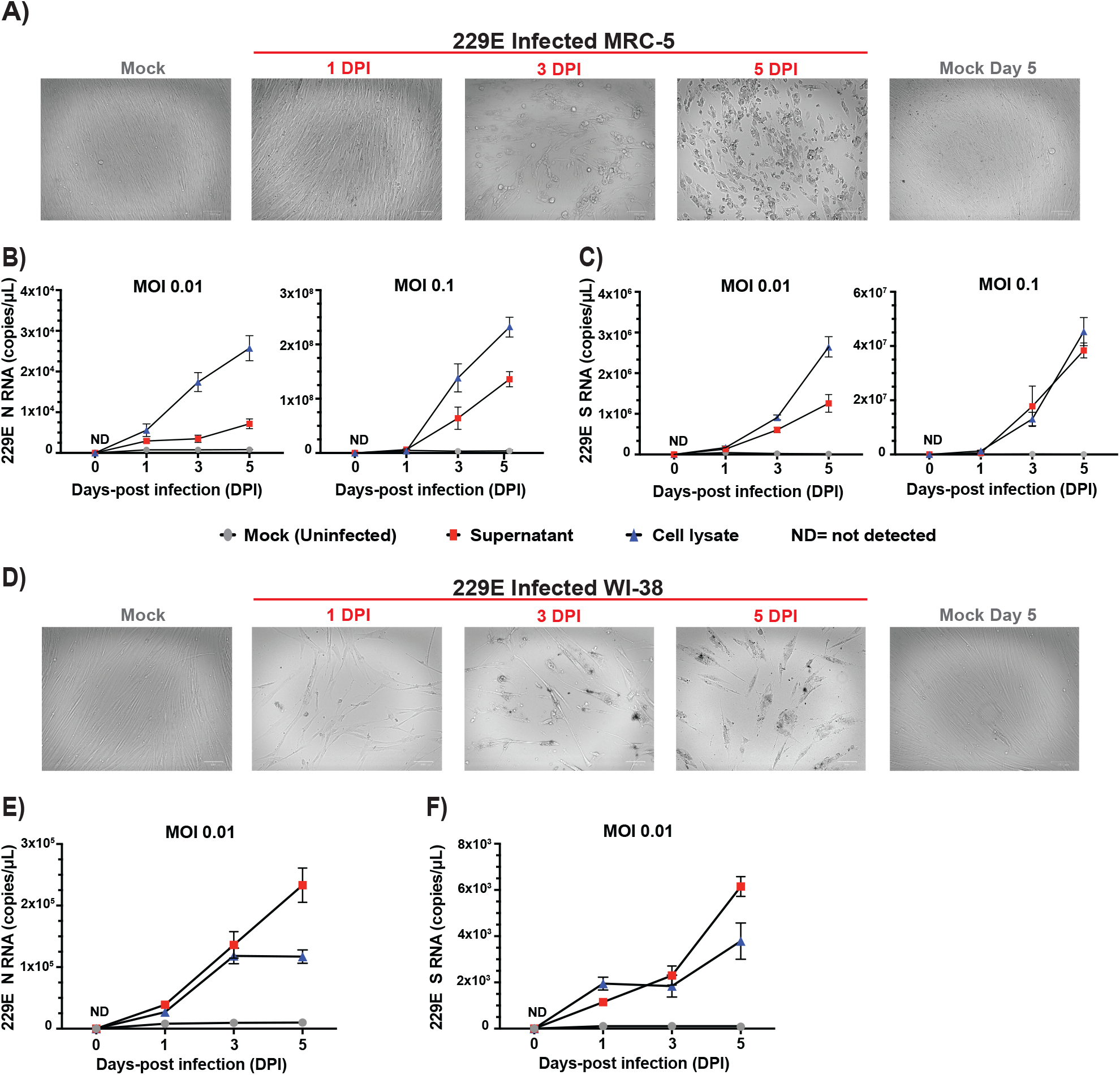
CPE evaluation and replication kinetics of 229E infection. (A) MRC-5 cells or WI-38 cells (D) were infected with HCoV-229E (virus titer: 5.64 ×10^6^ TCID50/ml) at a MOI of 0.01. CPE of the infected cells along with mock (uninfected cells) were monitored for morphological changes in the cell monolayer. Images of the cells were captured at different days post infection (day 0 to day 5 dpi) along with uninfected mock by ZOE Cell Imager. The white reference bar is 100 microns. MRC-5 cells (B and C) or WI-38 cells (E and F) were infected with 229E virus (MOI= 0.01 and/or 0.1) and cell lysates and supernatants were harvested at 0, 1, 3 and 5 days post-infection (dpi). Viral RNA copies of nucleocapsid (N), panels B and E, and spike (S), panels C and F, were then quantified by ddPCR. ND; not detected. Data represents 3 biological replicates with 3 technical replicates per time point (n = 9).

**Figure 4.**
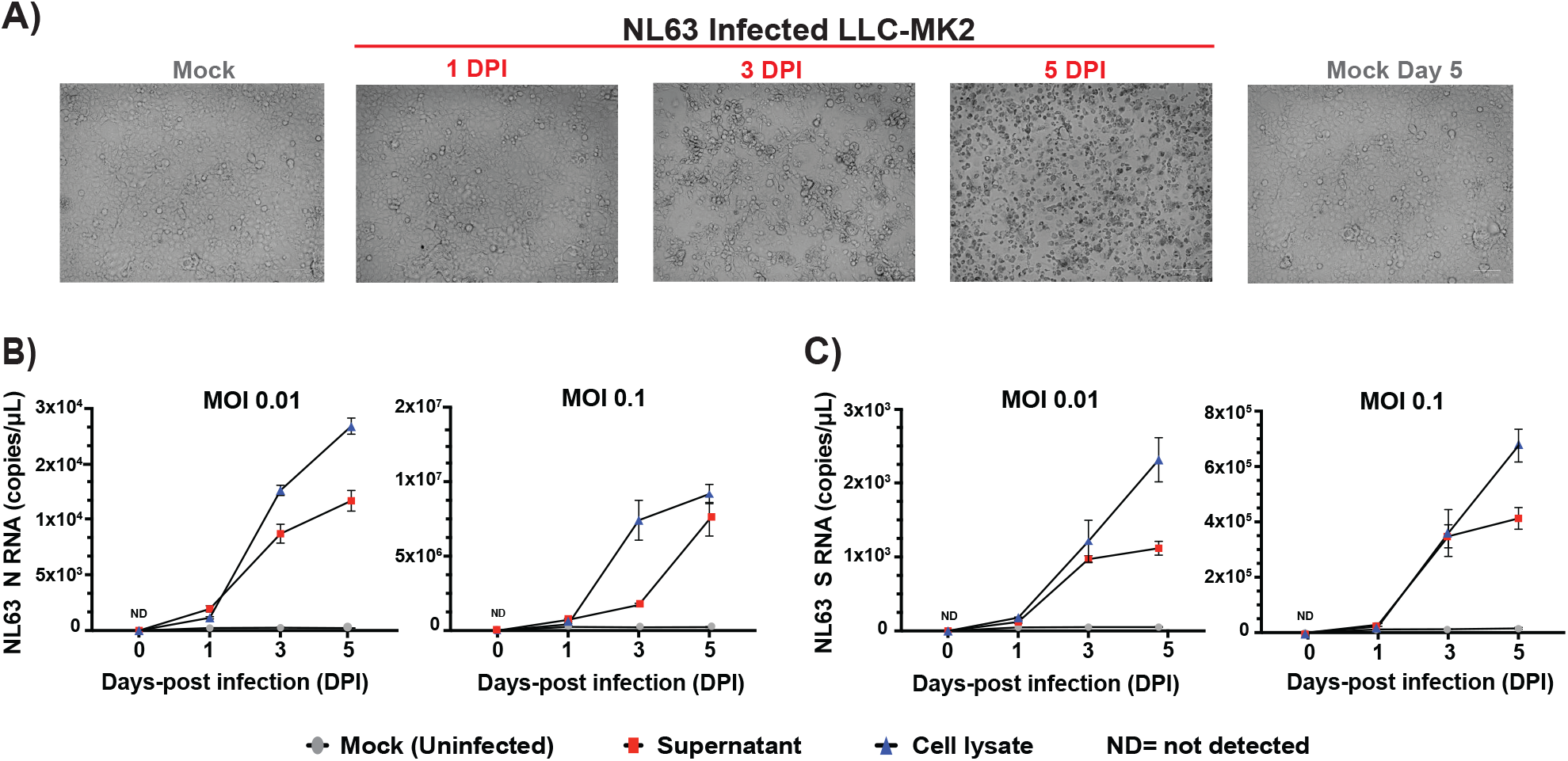
CPE evaluation and replication kinetics of NL63 infection. (A) LLC-MK2 cells infected with NL63 (virus titer: 1.58 ×10^6^ TCID50/ml) at a MOI of 0.01. CPE of the infected cells along with mock (uninfected cells) were monitored for morphological changes in the cell monolayer. Images of the cells were captured at different days post infection (day 0 to day 5 dpi) along with uninfected mock by ZOE Cell Imager. The white reference bar is 100 microns. LLC-MK2 cells were infected with NL63 virus (MOI= 0.01 and/or 0.1) and cell lysates and supernatants were harvested at 0, 1, 3 and 5 days post-infection (dpi). Viral RNA copies of nucleocapsid (N), panel B, and spike (S), panel C, were then quantified by ddPCR. ND; not detected. Data represents 3 biological replicates with 3 technical replicates per time point (n = 9).

### Characterization of OC43, 229E and NL63 viral replication kinetics

The culture supernatants and the cell lysates of permissive cells were infected with OC43, 229E and NL63 at a MOI of 0.01 and were harvested at days 1, 3 and 5 post infection. For OC43 with MRC-5 and WI-38 (**Fig. 2B-C, 2E-F, 2H-I**), 229E (**Fig. 3B-C, 3E-3F**), and NL63 (**Fig. 4B-C**) cell lysates show similar or higher vRNA copies of N and S in comparison to supernatants. However, for OC43 propagation in HCT-8 cells, we observe higher vRNA copies of N and S in the culture supernatant than in the cell lysate (**Fig. 2B and 2C**). We also see in these cells an increase in OC43 N and S vRNA copies starting from day 3 to day 5 post infection and a drop in S vRNA levels in the lysates for HCT-8 at day 5 pi (**Fig. 2B and 2C**), likely due to advanced cell death. Considering data from previous reports of NL63 infection in LLC-MK2 cells, the lower NL63 N and S vRNA levels detected in the supernatants compared to the lysates may be due to inefficient release of virions in the supernatants (17, 29, 35). Also, total vRNA copies detected varied depending on the cell types tested. For OC43 and 229E, WI-38 appeared the most favorable cell line for vRNA N expression, with ranges above 10^6^ copies/μL for OC43 and above 10^5^ copies/μL for 229E in the lysates at day 7 (**Fig. 2E and 3E**). Additionally, for all conditions tested, except infection of MRC-5 by 229E (**Fig. 3B and 3C**) and LLC-MK2 infection by NL63 (**Fig. 4B and 4C**), N vRNA levels are consistently several logs higher than S levels. For these combinations of virus and cells it was the opposite where S transcripts were more abundant than N. Overall, the results show that HCoV display variable replication kinetics depending on the specific virus and permissive host cell combination.

### Cell surface marker expression on HCT-8 cells following OC43 infection

Here, we characterized surface expression of GD3, CD13, CD147, CD326 and ACE2 protein on HCT-8 prior and after infection with OC43. CD13 (aminopeptidase N; APN) is the main surface protein receptor for 229E infection and is highly expressed on respiratory epithelial cells (12, 36). ACE2 is the primary receptor for NL63, and SARS-CoV and SARS-CoV-2 (16, 37, 38). CD147 is a transmembrane glycoprotein expressed on epithelial and immune cells that is proposed to be an alternative receptor for SARS-CoV and SARS-CoV-2 infection (39, 40). It is unknown if it plays a role in NL63 entry. CD326 is a marker of colon carcinoma cells that was used here as a control, it is known to be highly expressed on HCT-8 cells. OC43 binds to 9-*O*-acetylated sialic acid attached to oligosaccharide molecules located on glycoproteins on the surface of cells (14, 41). As such, this virus does not have a known specific protein surface receptor. However, ganglioside GD3 (9-*O*-acetyl-GD3) can be used by OC43 as a cellular entry receptor and is highly expressed in human leukemia cells (42).

Our assays detected low expression (8%) of GD3 in uninfected HCT-8 cells **(Fig. 5A and 5D).** We present the data as a scatter plot of a single representative assay **(Fig. 5A-5B)** and the compiled data summary of two-three biological repeats **(Fig. 5D and 5E).** Histogram plots of stained uninfected and infected cells relative to the corresponding isotype controls: IgG1 (CD13 and CD147) and IgG2b (CD326) and goat IgG (ACE2) are presented in **Figure 5C**. After infection of HCT-8, significantly fewer cells expressed the OC43 attachment receptor GD3 (2% of total) and those that did, expressed less of it as shown by a reduction of about 2-fold in mean fluorescence intensity (MFI) (**Fig. 5D and 5E**). We also detected expression of CD147 (100%) and CD326 (100%) surface antigens in all cells, and lower levels of CD13 (12%) and ACE-2 (17%) expression (**Fig. 5A and 5D**). Interestingly, infection with OC43 lead to a significant upregulation of ACE2 expression (30%) when compared to uninfected cells **(Fig. 5D and 5E)**. In contrast, we observed significant downregulation of CD326 MFI following infection (**Fig. 5E**), while there were no differences in expression levels detected for CD326, CD13 and CD147 (**Fig. 5D**). We did not present surface marker expression on WI-38 cells infected with 229E or OC43, or MRC-5 infected with OC43 given the intense and rapid CPE and cell death that compromised flow cytometry analysis.

**Figure 5.**
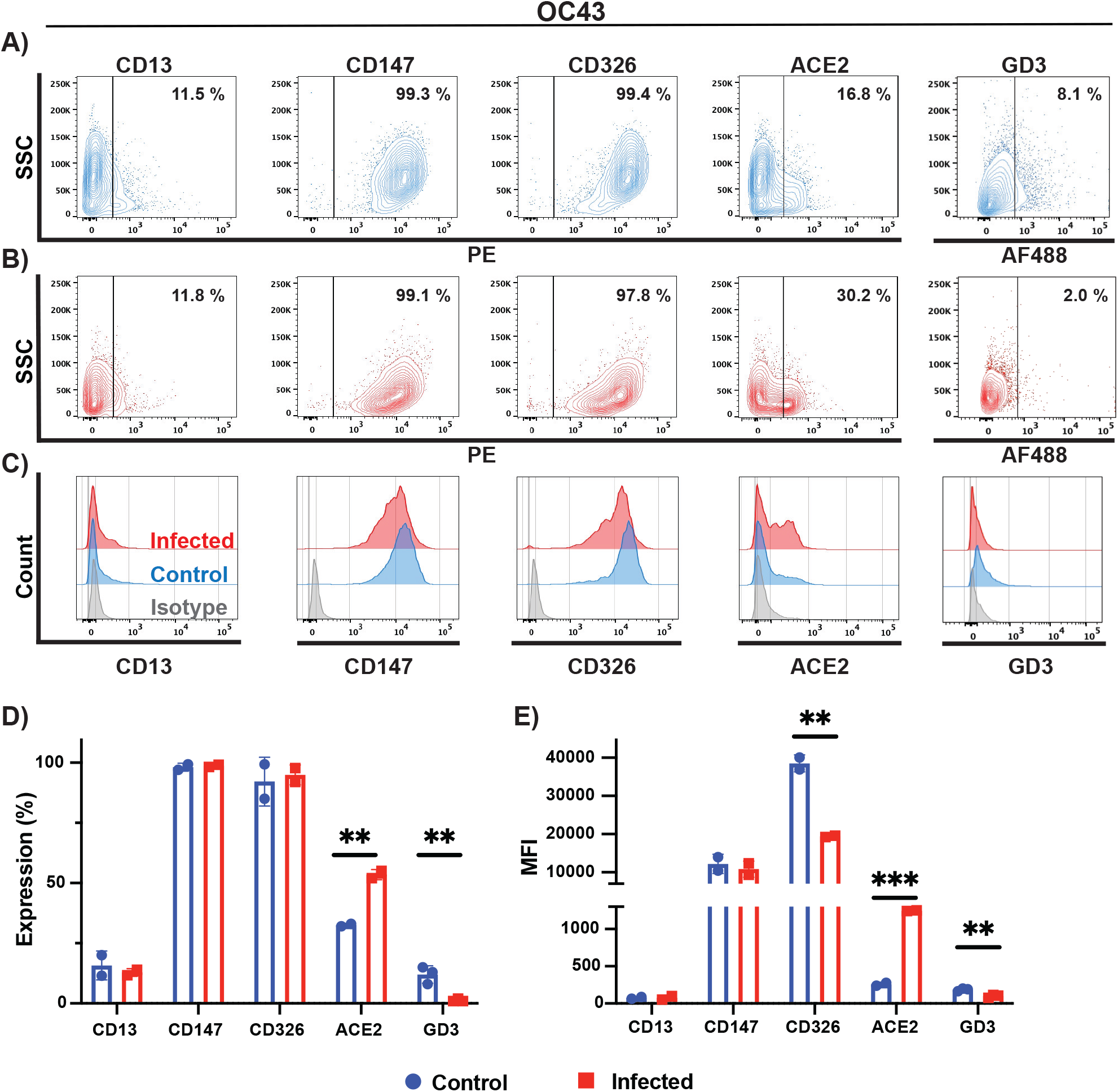
Phenotypic characterization of cell surface markers of OC43 infected cells. Phenotypic characterization of (A), uninfected (blue), and (B), OC43 infected HCT-8 cells (red). CD13, CD147, CD326, ACE2 and GD3 expression were probed by flow cytometry. (C) Histogram plots of control and infected HCT-8 cells relative to isotype control (gray). Histograms plots showing (D), percentage of cells expressing a given marker, and (E), mean fluorescence intensity (MFI). Data in (D) and (E) represent the mean of two biological replicates. Statistical analysis was performed with the multiple unpaired t-test with Welch’s correction. Only results that reached statistical significance were identified: * *p* ≤ 0.033, ** *p* ≤ 0.002, *** *p* ≤ 0.0001, **** *p* ≤ 0.0001.

### Cell surface marker expression on MRC-5 cells following 229E infection

Next, we characterized surface expression of CD13, CD147, CD326 and ACE2 proteins in uninfected and infected MRC-5 cells with 229E. No expression of GD3 was detected on MRC-5 or LLC-MK2 cells (data not shown). Nearly all MRC-5 cells expressed high levels of CD13 (91%) and CD147 (100%) surface antigens and much lower levels of CD326 (0.8%) and ACE2 (13%) markers (**Fig. 6A and 6C).** Infection of MRC-5 cells with 229E virus lead to an important decrease in cells expressing 229E cellular receptor CD13, from 99% to 55% (**Fig. 6A and 6B**). MFI and the percentage of cells expressing each marker prior and following infection with OC43 are summarized in **Figure 6D and 6E**. Although CD326 and ACE2 expression were slightly reduced following 229E infection, these did not attain statistical significance.

**Figure 6.**
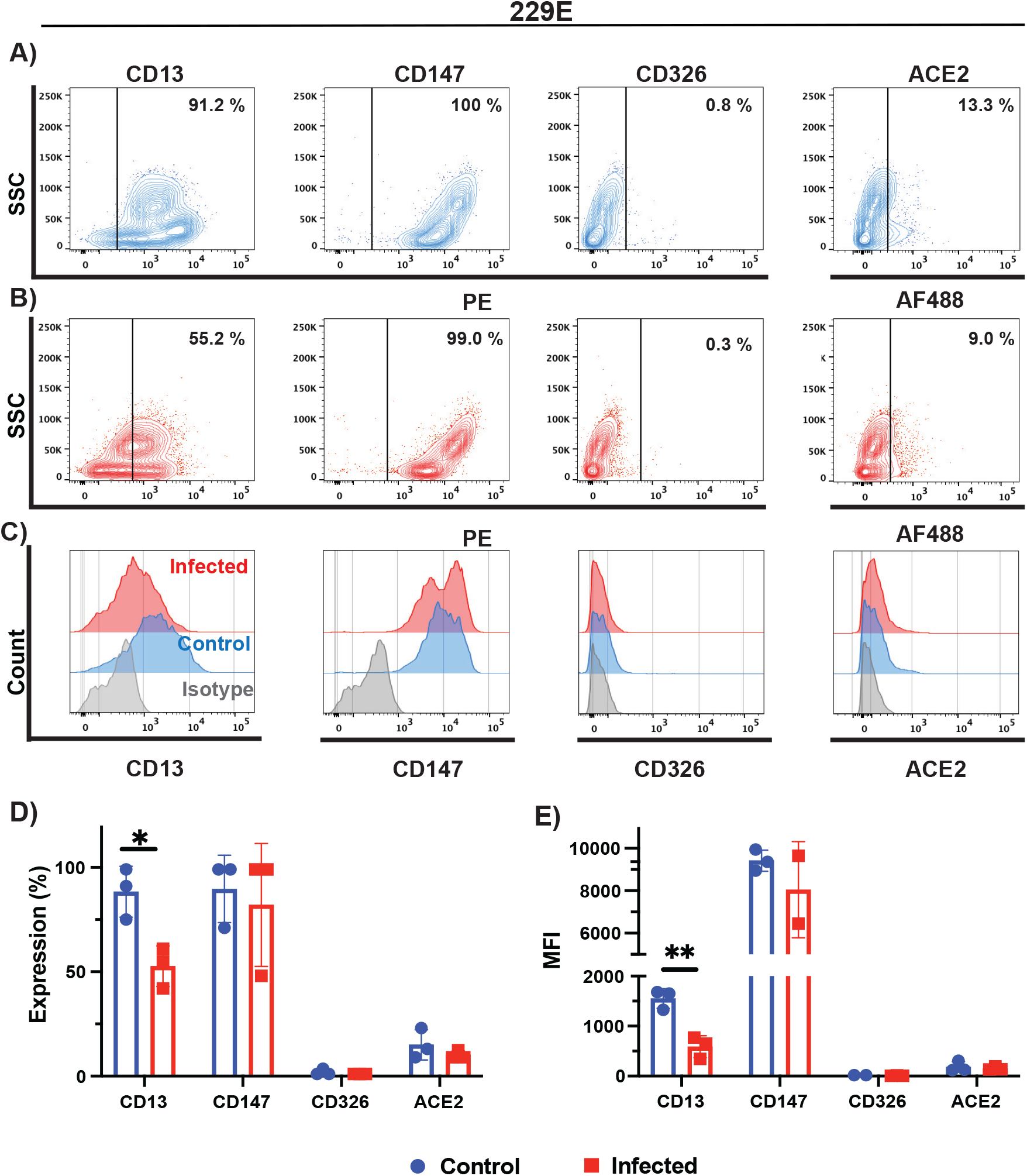
Phenotypic characterization of cell surface markers of 229E infected cells. Phenotypic characterization of (A), uninfected (blue), and (B), 229E infected MRC-5 cells (red). CD13, CD147, CD326, and ACE2 expression were probed by flow cytometry. (C) Histogram plots of control and infected MRC-5 cells relative to isotype control (gray). Histograms plots showing (D), percentage of cells expressing a given marker, and (E), mean fluorescence intensity (MFI). Data in (D) and (E) represent the mean of two biological replicates. Statistical analysis was performed with the multiple unpaired t-test with Welch’s correction. Only results that reached statistical significance were identified: * *p* ≤ 0.033, ** *p* ≤ 0.002, *** *p* ≤ 0.0001, **** *p* ≤ 0.0001.

### Cell surface marker expression on LLC-MK2 cells following NL63 infection

Lastly, we characterized surface expression of CD13, CD147, CD326 and ACE2 proteins in uninfected and infected LLC-MK2 cells with NL63 virus. We detected high expression of CD13 (81%) and CD147 (99 %), and lower expression of ACE2 (27%) surface receptor in uninfected LLC-MK2 cells **(Fig. 7A)**. Infection with NL63 lead to a 2.3-fold increase in cells expressing NL63 cellular receptor ACE2 (**Fig. 7B-D).** Although we did observe a small upregulation of CD13 and CD326 expression in infected cells, these did not reach statistical significance (**Fig. 7D**). Previous studies have shown the involvement of CD147 in SARS-CoV-2 infections (40, 43). However, CD147 expression remained unaffected by NL63 infection as shown here **(Fig. 7D and 7E)**.

**Figure 7.**
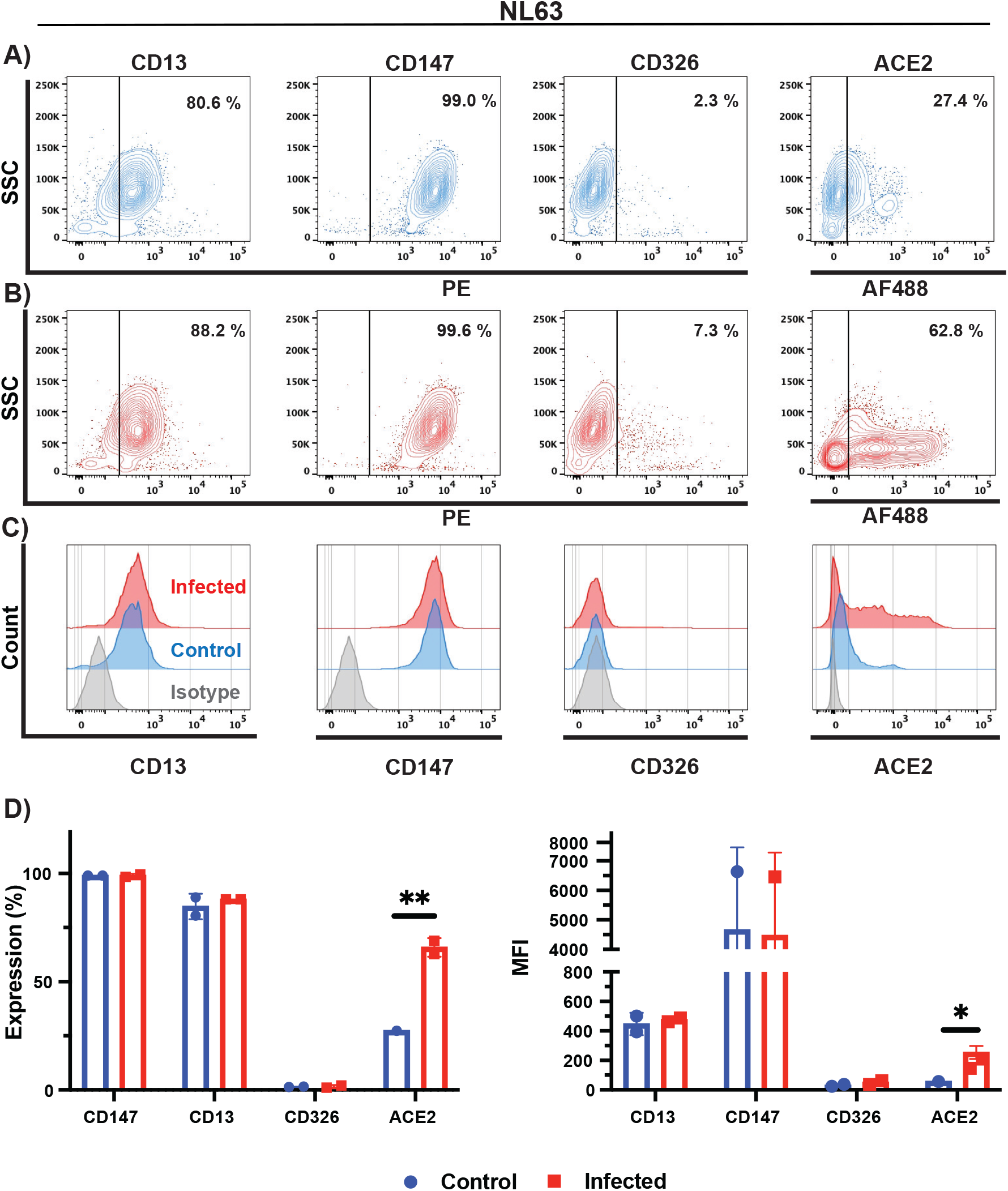
Phenotypic characterization of cell surface markers of NL63 infected cells. Phenotypic characterization of (A), uninfected (blue), and (B), NL63 infected LLC-MK2 cells (red). CD13, CD147, CD326, and ACE2 expression were probed by flow cytometry. (C) Histogram plots of control and infected LLC-MK2 cells relative to isotype control (gray). Histograms plots showing (D), percentage of cells expressing a given marker, and (E), mean fluorescence intensity (MFI). Data in (D) and (E) represent the mean of two biological replicates. Statistical analysis was performed with the multiple unpaired t-test with Welch’s correction. Only results that reached statistical significance were identified: * *p* ≤ 0.033, ** *p* ≤ 0.002, *** *p* ≤ 0.0001, **** *p* ≤ 0.0001.

### Soluble ACE2 protein and anti-ACE2 antibody effectively inhibits NL63 infection

Given that ACE2 is the major receptor for NL63 infection, and having now established viral infection assays to measure vRNA expression, we wanted to investigate the efficiency of blocking antibodies and soluble ACE2 (sACE2) as infection prevention strategies. While therapeutic approaches that target ACE2 have been developed for SARS-CoV-2, it is not known if these can also inhibit NL63 infection (44–49).

LLC-MK2 cells were infected with NL63 (MOI of 0.1) and supernatants were harvested 5 days later. Supernatants (MOI equivalent of 0.1) were pre-treated with sACE2 or control protein (soluble ovalbumin) for 30 min. prior to infection of LLC-MK2 cells (**Fig. 8A and 8C).** Similarly, LLC-MK2 cells were pre-treated with a blocking anti-ACE2 mAb or an unrelated control antibody (anti-DC-SIGN) for 30 min. prior to infection **(Fig. 8B and 8D)**. Five days after infection, we measured S and N vRNAs in the supernatants and lysates of infected cells. Both the culture supernatants and cell lysates of virus-infected LLC-MK2 cells with prior treatment of the virus with sACE2 led to a significantly attenuated production of N and S vRNA copies, of about 83% and 89% in supernatants and 96% and 88 % in lysates at the highest dose of 10 ³g/mL, respectively **(Fig. 8C).** Similarly, treatment of LLC-MK2 cells with the human monoclonal ACE2 (anti-ACE2 mAb) blocking antibody shown to be capable of SARS-CoV-2 inhibition (49) also inhibited NL63 infection in a dose-dependent manner **(Fig. 8B and 8D)**. Both the NL63 N and S vRNA levels were significantly reduced by 76% and 68 % in supernatants and 89% and 86 % in lysates at the highest dose of 120 mg/ml, respectively **(Fig. 8D).** However, neither soluble ACE2 nor anti-ACE2 were capable of completely blocking the infection in these conditions.

**Figure 8.**
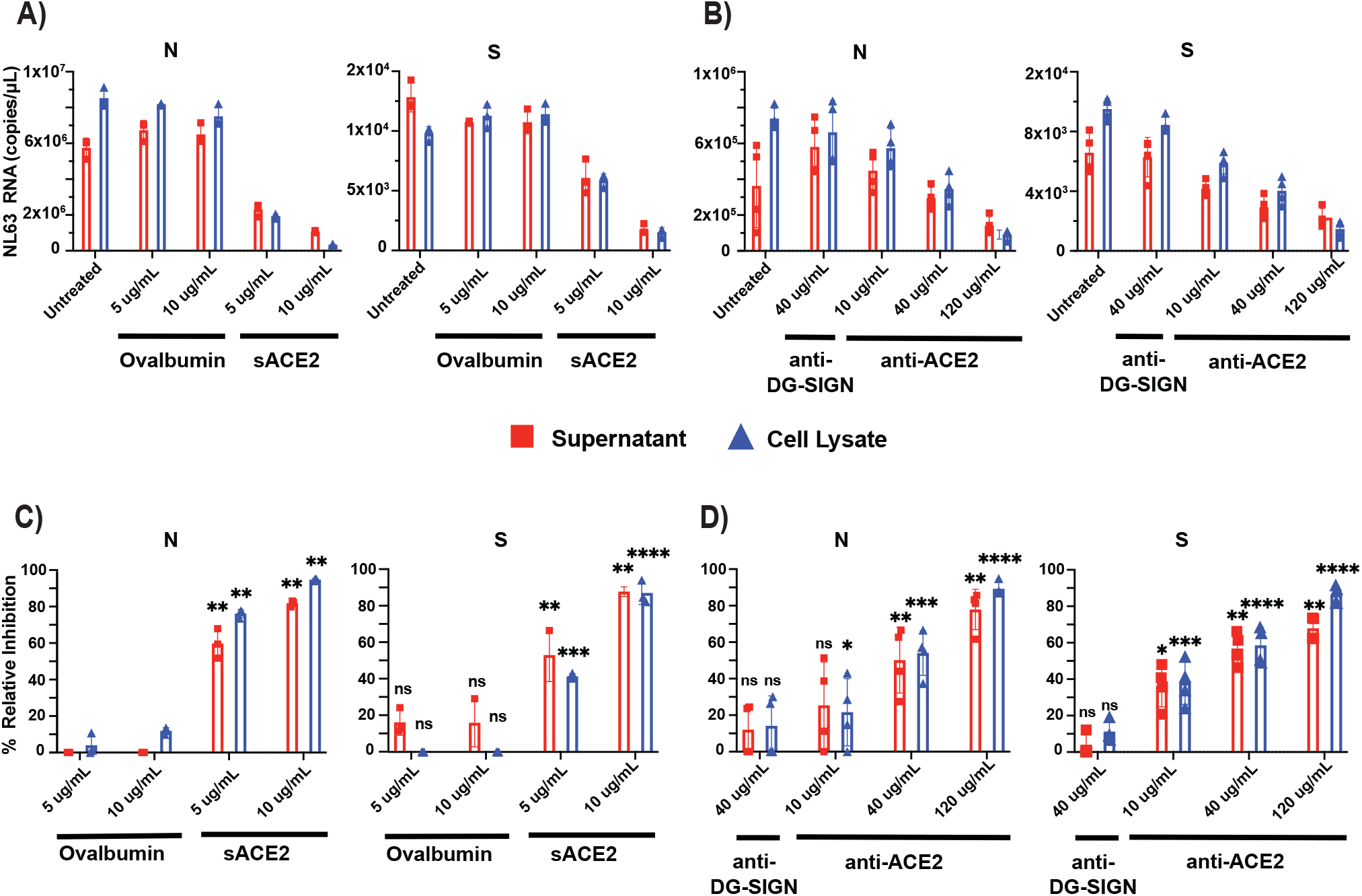
Soluble ACE2 protein and anti-ACE2 monoclonal blocking antibody inhibits NL63 infection in LLC-MK2 cells. (A) Treatment with soluble ACE2 (sACE2) or control protein ovalbumin at 5 and 10 mg/mL was pre-incubated with NL63 virus at MOI 0.1 (virus titer: 8.89 ×10^5^ TCID50/ml)) for one hour and the protein and virus mixture complex was added to the LLC-MK2 cells. (B) Similarly, cells were incubated for one hour with human anti-ACE2 monoclonal blocking antibody (10, 40 and 120 µg/ml), or unrelated control antibody (anti-DC-SIGN; 40 µg/ml) followed by addition of NL63 virus (MOI 0.1). Five days post infection, both culture supernatants and cell lysates were harvested for quantification of nucleocapsid (N) and spike (S) vRNA copies by ddPCR. Percent relative viral inhibition following treatment with C) sACE2 or D) anti-ACE2 monoclonal antibody compared to their respective untreated control. Data represents 2 biological replicates with 3 technical replicates per condition (n = 6). Statistical analysis was performed using an unpaired t-test with Welch’s correction (P<0.05) relative to untreated control. n.s denoted as no statistical difference; * *p* ≤ 0.033, ** *p* ≤ 0.002, *** *p* ≤ 0.0001, **** *p* ≤ 0.0001.

## DISCUSSION

An in-depth understanding of common cold seasonal coronavirus transmission, infection, pathogenicity, and the impact of reoccurring infections on human health is incomplete (2, 50, 51). In fact, only very recently was the functional receptor for HKU1 identified (15). Moreover, given that these viruses have zoonotic and co-infection potential to create new recombinant strains, they are of particular interest in light of the current SARS-CoV-2 pandemic (52, 53). This work studied the characteristics of HCoVs-host cell interactions, CPE and replication kinetics in commonly used human and animal cell lines. We used cell culture models of human lung, colon cancer and rhesus monkey kidney cell lines that are fibroblastic and epithelial in origin as platforms to study the impact of infection on the host cell and attachment receptors.

Viral replication and CPE analysis demonstrated that of all the cell lines tested (**Table 1),** MRC-5, HCT-8 and LLC MK2 are the most suitable in our conditions to study OC43, 229E and NL63 *in vitro*. While WI-38 did get infected by OC43 and 229E, the cells exhibited severe CPE that compromised our ability to perform flow cytometry analysis. We also considered to perform plaque assays for our study, however we were unsuccessful to obtain sufficiently defined plaques that were amenable to quantification following OC43, 229E and NL63 infection (data not shown). Cells that are permissive to infection do not necessarily demonstrate CPE or support plaque formation using standard methods (26, 30). However, some other groups have reported, albeit with varying success, plaques following HCoV infection using specialized protocols (4, 54). Viral RNA analyses revealed that OC43, 229E and NL63 replication in host cells steadily increased by day 3 pi, followed by CPE and dysmorphic cell appearances (**Fig. 1-4**). This is supported by previous reports of HCT-8 and MRC-5 infections with OC43 where the cells displayed obvious CPE at day 2 or 3 pi (26). However, the expression profiles and abundance of vRNAs in both supernatants and cell lysates for each virus-host cell pair varied significantly over the 5-day analysis period. Accumulation of vRNA was consistently higher in cell lysates than in supernatants, except for OC43 infecting HCT-8 cells where it was the opposite (**Fig. 2B and 2C**). Also, there was a general trend whereby more N than S vRNA was measured in both lysates and supernatants for all virus-host cell pairs except for MRC-5 cells infected by 229E that displayed 100-fold higher levels of S than N (10^4^ vs 10^6^ copies per mL) (**Fig. 3B and 3C**). The overall canvas that emerges is one where cell-intrinsic factors influence the outcome of viral replication and release. These can include restriction factors that inhibit viral, for instance, budding and egress, or hinder viral genome replication and stability such as RNA-editing enzymes (55–58). Interferon and antiviral responses may also play an important role in viral spread and CPE in cell cultures (19).

Some viruses modulate surface receptors and co-receptors on the surface of the cell. For example, in the case of HIV, the virus downregulates CD4 expression on lymphocytes via the viral nef protein to avoid co-infections and superinfections that could lead to host cell genome instability (59). In this study, we analyzed the surface expression of CD13, CD147, CD326, GD3 and ACE2 on infected and uninfected MRC-5, HCT-8 and LLC-MK2 cells by flow cytometry. We tracked both the proportion of cells expressing the receptor and the receptor expression intensity, MFI. While we observed a 4-fold decrease in GD3 on HCT-8 cells following OC43 infection, as well as a 1.7-fold decrease in CD13 expression following 229E infection, we were surprised to observe a significant increase in ACE2 receptor expression and MFI following NL63 infection of LLC-MK2 cells (2.3-fold), but also following OC43 infection (1.8-fold). Induction of ACE2 expression following infection with NL63 and the highly prevalent OC43 may be a source of concern for coronavirus co-infections. Although cytokine and interferon responses induced during the primary infection by OC43 and NL63 will likely have an inhibitory effect on co-infection by another coronavirus utilizing ACE2 for entry (60), a strong induction in receptor expression could tilt the balance in favor of the incoming virus. There is currently very little information in the published literature about the frequency of such co-infections and their possible role in creating recombination events with SARS-CoV-2 (61).

Currently, there are no approved or licenced antiviral compounds for the treatment of the common cold caused by seasonal HCoV infections. Previous studies have shown that sACE2 or engineered variants of sACE2 may be beneficial for the treatment of COVID-19 infection (18, 62). LLC-MK2 cells treated with the highest dose of sACE2 (120 µg/mL) reduced the expression of N and S vRNA of NL63 by 90% in viral lysates (**Fig. 8B and 8D)**. Similarly, human anti-ACE2 monoclonal blocking antibody inhibited in a dose-dependent manner (10, 40 and 120 µg) NL63 infection (**Fig. 8B and 8D**). These data indicate that targeting the entry receptor appears to be very effective at inhibiting this virus. However, we also attempted to use a polyclonal anti-ACE2 antibody to block NL63 without success (data not shown), suggesting that there may be few critical neutralizing epitopes in the spike-ACE2 interface for NL63 entry and that these should be prioritized to develop a therapy. Soluble ACE2 also proved to a capable inhibitor of NL63, but to somewhat a lesser degree.

In summary, our study showed that HCoVs OC43, 229E and NL63 display differential replication kinetics and CPE in permissive host cells highlighting the underlying involvement of cell-intrinsic factors that influence the expression of viral genes and virus egress. We also demonstrate that HCoV OC43 and NL63 infection upregulate the expression of the entry receptor ACE2 on host cells, raising opportunities for possible co-infection and recombination with other HCoVs (61, 63, 64). How HCoVs recombine and evolve in a given host remains an important topic to better the understand the emergence of new viral strains of coronaviruses.

## ACKNOWLEDGEMENTS

This study was supported by a COVID-19 Rapid Response grant by the Canadian Institutes of Health Research (CIHR; #VR2-172722) to M-A Langlois. M.M. holds an Ontario Graduate Scholarship. Y.G. holds a Canadian Institute of Health Research (CIHR) Frederick Banting and Charles Best graduate scholarship (CGS-Doctoral, 476885). N.C holds a CIHR Frederick Banting and Charles Best graduate scholarship (CGS-M). M.-A.L. holds a Faculty of Medicine Chair of Excellence in Pandemic Viruses and Preparedness Research.

## CONFLICTS OF INTERESTS

The authors declare no conflict of interest relevant to the present manuscript. The funding sources played no role in the study design, data collection, analysis, writing of the manuscript, and the decision to publish.

### Data availability statement

The data used in this study can be obtained by request from M-A.L.

### Author contributions

V.S., M.M. and M.-A.L. designed the study. V.S., M.M., N.C., M.S., J.H.S. and S.D. performed the experiments. V.S., M.M., Y.G. and M.-A.L. analyzed the data. V.S., M.M., Y.G. and M.A.L wrote the manuscript. All authors edited the final version of the manuscript.

